# The Size of Fields in Biomedical Sciences

**DOI:** 10.1101/2023.06.13.544650

**Authors:** Quigly Dragotakes, Arturo Casadevall

## Abstract

Scientific research output has increased exponentially over the past few decades, but not equally across all fields of study, and we lack clear methods for estimating the size of any given field of research. Understanding how fields grow, change, and are organized is essential to understanding how human resources are allocated to the investigation of scientific problems. In this study we estimated the size of certain biomedical fields from the number of unique author names appearing in field relevant publications in the PubMed database. Focusing on microbiology, where the size of fields is often associated with those who work on a particular microbe, we find large differences in the size of its subfields. We found that plotting the number of unique investigators as a function of time can show changes consistent with growing or shrinking fields. We envision using unique author count to measure the strength of a workforce in any given field, analyze the overlap of workforce between fields, and compare how workforce correlates to available research funds and public health burden of a field.

## Introduction

As the knowledge domain of science increased there was a need for specialization^1^. Early scientists were generalists who could be expected to have some knowledge of the totality of human epistemic endeavors and were known as natural philosophers. For example, Newton created the calculus, formulated the laws of Newtonian physics, knew astronomy, was interested in theology, and may have spent much of his time working on alchemy, a precursor discipline to chemistry. However, by the 19^th^ century the knowledge base had grown so large that scientists specialized in the different areas of the natural sciences such as physics, chemistry, and biology. These in turn underwent further specialization in the 20^th^ century into discrete fields that focused on specific problems.

Today, science is organized into fields and many fields have several subfields. We will use the fields of microbiology and immunology as examples, as these are the ones we are most familiar with. Both microbiology and immunology are subfields of the larger field of biomedical sciences, which is in turn a subfield of biology^1,2^. Hence, fields constitute subgroupings of scientists working on discrete problems often within fussy epistemic boundaries. Fields are also the sociological units by which science is organized since they constitute a major source of friends and personal contacts for scientists^3^. Herrera et al used a network analysis to map the connectivity and size among fields within Physics^3^. Chevalarias and Cointet developed a method to infer phylomemetic patterns from the published literature and used its density to ascertain the growth and decline of fields^4^. The question of field size is important for understanding how human resources are invested in areas of science and could be significant for the scientific progress given that large fields may stymie the development of new ideas and concepts^5^.

Despite the importance of field organization to science we could find little or no information on the size of fields. Although scientists anecdotally know that some fields are larger than others, remarkably little has been done to quantitate the size the fields. This is an important problem because the size of a field is a measure of how much human capital is devoted to a particular problem. For example, when the COVID-19 pandemic began in 2019 it may have been useful to know the number of scientists with experience in coronavirus or viral vaccines as this would have provided a measure of human resources available to confront such a threat. Knowing the size of fields is also important when considering the allocation of the efficient allocation of scarce resources. However, estimating the size of fields is not an easy task. Scientists move between fields and often engage in cross-field research, creating fuzzy boundaries that defy easy categorization. Assuming one can identify a measure for field size there are other obstacles to accurate enumeration. For example, many scientists published in this journal work on various problems simultaneously and therefore may belong to more than one field at any given time. In this regard, the work of these authors could fit within the fields of immunology or microbiology, or their interface, depending on how their contributions are assessed. Second, fields evolve with time with some increasing in size and others shrinking. In this study we approach the problem of size of fields using bibliometric approaches whereby the number of scientists with a given name is associated with a subject and have developed software that allows one to estimate the size of fields. We use microbiology to explore this topic since this is subfield of biomedical sciences that is sectioned by the microbes studied^2^.

We hypothesize that the size of a given field can be estimated by counting the total unique authors in published articles of said field. In this study, we estimated the size of subfields within the field of microbiology by counting the number of unique names identified with specific microbes and find large variation. We anticipate that the number of papers associated with a topic is also a measure of the size of a field but predict that counting authors rather than papers will provide a better estimate of field size since each individual is unique and fields are composed of people, not papers. For example, a publication describing a microbe interacting with a macrophage could belong to the fields of microbiology, immunology, and cell biology while an author in the paper is a unique entity who is potentially traceable by name or ORDIC. Furthermore, focusing on individuals mitigates confounders arising from differences in laboratory productivity. For example, a field composed 100 laboratories that each produce one paper per year is larger than a field composed of 1 laboratory that produces 100 papers per year, but this distinction could not be made by only counting research output.

## Methods

A given field was denoted by a search term, which was used to query the PubMed database (https://pubmed.ncbi.nlm.nih.gov/). All found publications were then downloaded using Entrez Direct and a list of all authors was taken from the recorded author bylines^6^. Unique names were determined by the first and last names documented in the PubMed database. Data was only collected for articles with a publication date and authors with a recorded first and last name entry. Funding information was collected from NIH RePORTER (https://reporter.nih.gov/) with search terms matching those of PubMed queries. Case burden information was gathered from literature and CDC national reporting^7,8^. Author information was parsed and analyzed in R (version 4.3.0)^9^.

## Results

### Size of a Field can be Estimated by Counting Unique Authors

We first estimated the size of given fields by counting all unique authors which appear in PubMed deposited articles. We found that each of these fields had a general upward trend in the number of unique authors over time, and that individual species searches heavily resembled that of the total genus searches (Figure 1A). We next investigated how closely the number of unique authors in a field correlated to the number of papers published in any given year. We found that there is a strong overall correlation between authors and publications, with the strength of the correlation in individual fields showing more variation (Figure 1B, Supplemental Figure 1). While the number of unique authors in a field was clearly reliant on publications being generated in said field, we can generate a deeper image of a field’s growth with authors. Counting authors accounts for trainees, collaborations, etc. which would otherwise be missed by just counting the publication itself.

**Figure 1.**
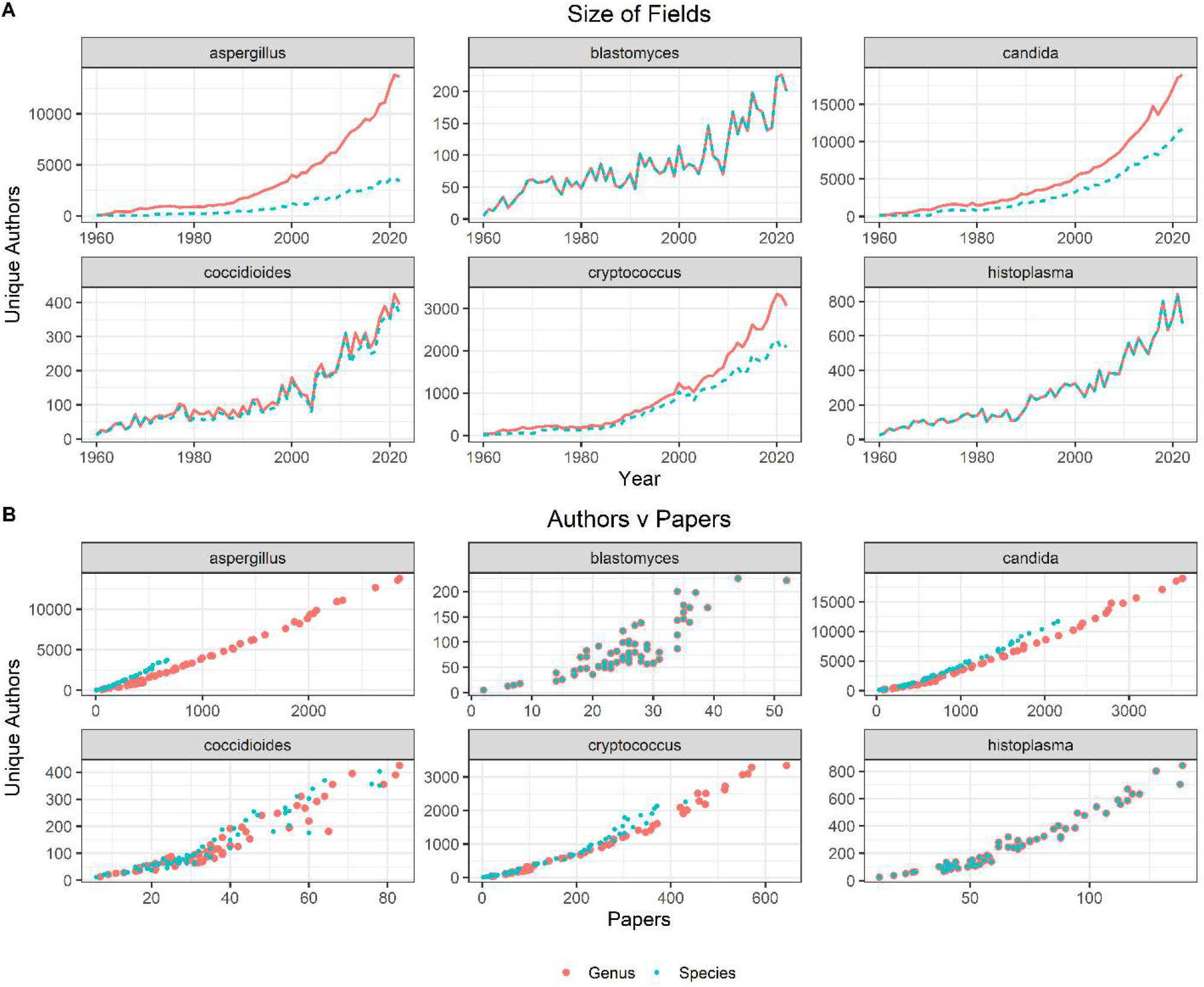
Estimating the Size of Fields by Author Count. **A**. The size of six significant fungal fields was calculated by searching PubMed for either just the genus (blue) name, or the genus and species (red) name. **B**. The number of unique authors in each field each year were compared to the total number of publications. An overall positive correlation was observed but varied by field. Species designations are as follows: *Aspergillus fumigatus, Blastomyces dermatitidis, Candida albicans, Coccidioides immitis, Cryptococcus neoformans*, and *Histoplasma capsulatum*.

### Field Size Correlates to Disease Burden

The number of cases attributed to a particular disease is a measure of the importance of that disease to society. Knowing the size of the fields working on specific diseases is a measure of the resources invested in studying a disease, which provides a mechanism for evaluating the consistency of resource allocations. We first investigated whether the total amount of funding for a given pathogen correlated to the size of a given field. We found that, generally, the increase in funding generally outpaced the increase in authors for all six of our fungal fields, resulting in higher dollar per author funding to the fields (Figure 2A). The trend remains largely the same after adjusting for inflation.

**Figure 2.**
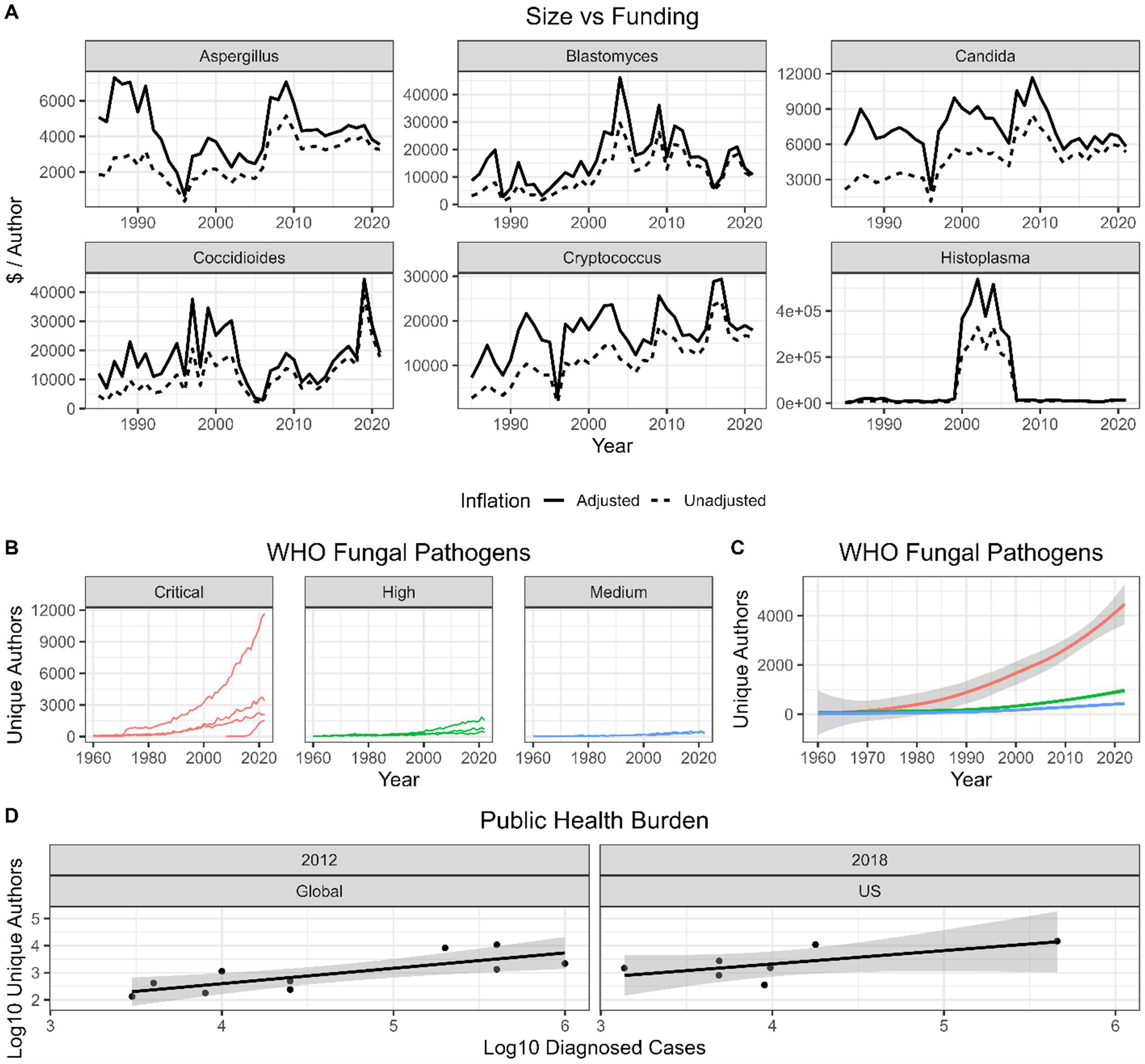
NIH Funding and Health Burden According to Field Size. **A**. We report the number of US Dollars ($) awarded to a field compared to the total authors within that field over the course of all available NIH Reporter data and adjusted for inflation according to average annual CPI for all urban consumers. **B-C**. Size of fields compared to global health burden of pathogen. We determined the size of field for each of the fungal pathogens recently categorized by the WHO in terms of criticality. Critical and High priority fungal fields are larger than medium, but all categories are growing steadily in size. **B**. A combined regression for unique author analysis of all fungal diseases of a given category, pooled together. **C**. Individual author analyses for each fungal pathogen within a category. **D**. Workforce compared to public health burden of given diseases. In both Global and US specific diseases the size of a field correlates to disease burden.

Scientists mostly study pathogens and diseases of interest to them and for which resources are allocated, so we next investigated whether fields naturally stabilize to sizes reflecting the case burden of the disease. The WHO recently released a list of fungal pathogen priorities, and we compared field size to the relative priority of each fungal disease. As expected, a larger proportion of the workforce is committed to critical priority pathogens, though the high priority field is experiencing rapid growth (Figure 2B-C).

To investigate more closely, we compared the size of fields working on various fungal diseases to the burden of disease caused by that disease. Unfortunately, current reporting of fungal diseases is limited and not standardized. However, we were able to obtain reliable estimates of various fungal disease diagnoses for two years and compare case burden to size of fields. We found that, in both years, number of authors reporting on a field correlated positively with case burden (Figure 2D).

We next investigated whether these trends held true outside of fungal diseases by expanding our search to 10 major human diseases. We observed largely similar results to the fungal field where workforce and funding was generally well distributed and correlated to case burden (Figure 3). Several spikes in funding are noticeable after major events involving particular diseases: the 2018 Salmonellosis outbreak, the 2016 Zika outbreak, etc. For the most part however, the workforce attributed to each category has been steadily increasing. The largest effects on workforce, as one would expect, seem to follow diseases uncommon or previously unseen to the US breaking out (Zika outbreak in 2016, novel Coronavirus in 2019, etc.)

**Figure 3.**
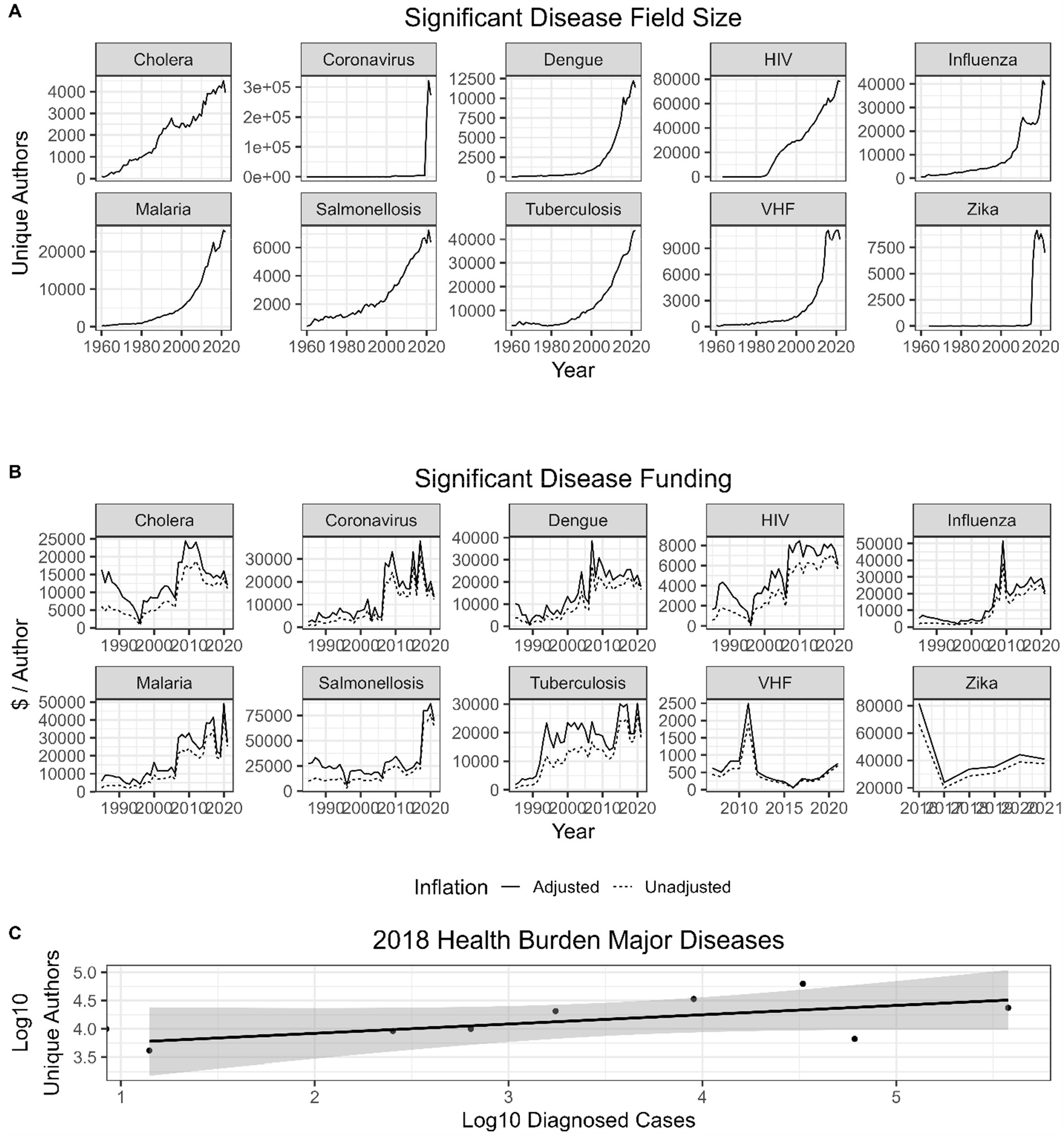
Size and funding of major human diseases. **A**. Size of major human disease fields counted by unique authors. **B**. Funding per author for significant human disease fields adjusted and unadjusted for inflation. **C**. Size of major disease fields compared to case burden in the US during 2018.

### Unique Authors Mirrors Activity and Use of Model Organisms and Methods

One would expect the number of unique authors in a field to mirror a specific model organism or pathogens use and outdated or early versions of methods are less likely to be used by new investigators. Thus, we analyzed several model organisms (*Saccharomyces cerevisiae, Danio rerio, Drosophila melanogaster*, and *Caenorhabditis elegans*), three pathogens we would not expect to continue increasing in size given advances in virology and vaccines (SV40, poliovirus, *Variola major)*, a field which experienced multiple popularity spikes from discoveries decades apart (phage), and three recent gene editing methods with overlapping userbases: zinc finger nucleases (ZFNs), transcription activator-like effector nucleases (TALENs), and clustered regularly interspaced short palindromic repeats (CRISPR). We observed expected patterns of field size according to use and prevalence over time for each of these categories (Figure 4).

**Figure 4.**
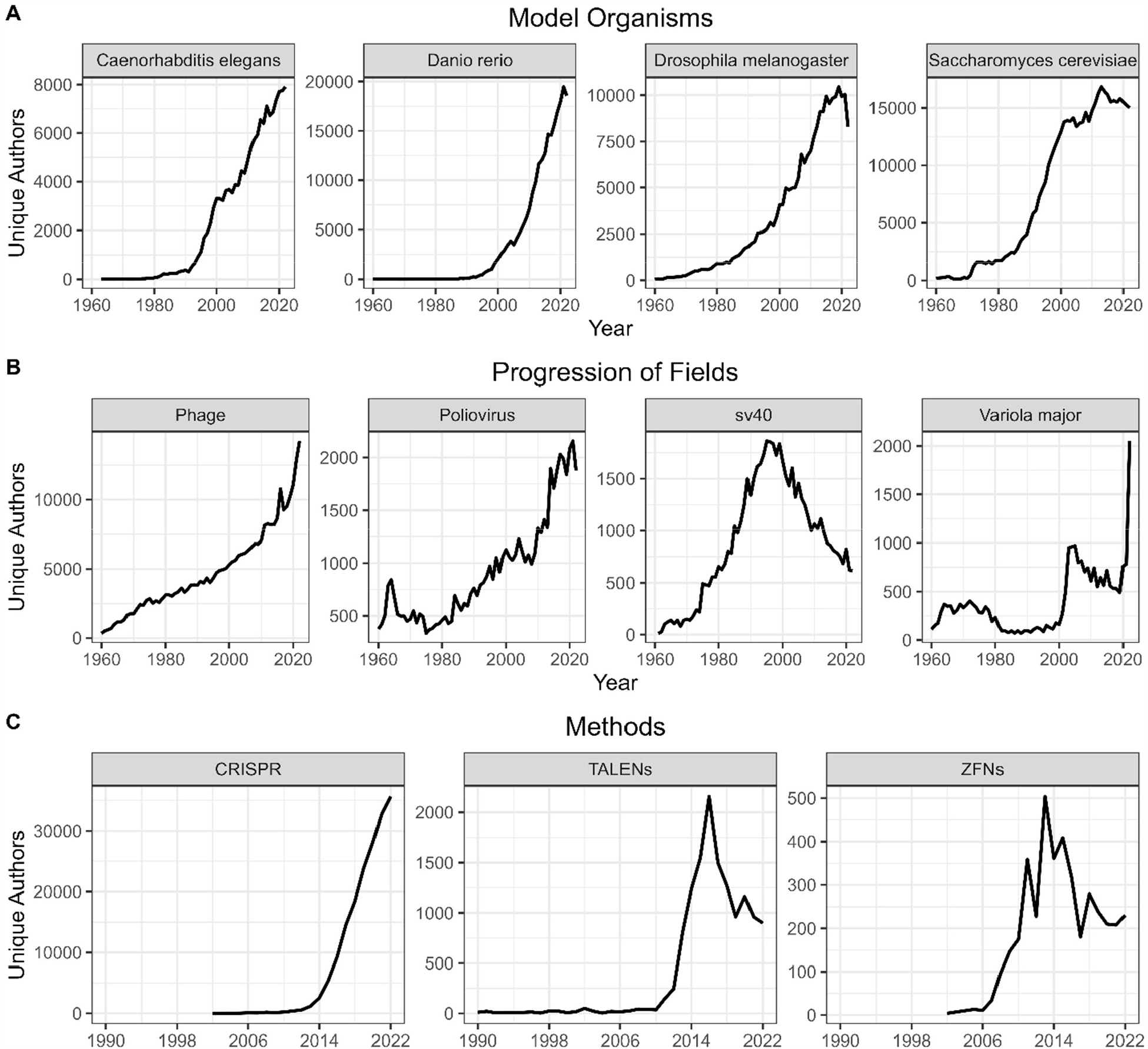
Growth and Shrinking of Fields. We analyzed four model organisms, four fields of research, and three iterations of gene editing technology to investigate how unique author count correlates to fields growing and changing over time. **A**. We found that the size of model organism fields correlated well with their increased use. Interestingly the Drosophila field eventually plateaued and was supplanted by the zebrafish. **B**. We found the authors in the SV40 field correlated well with its use in generating cancer models. Polio and smallpox mostly remained steady at lower numbers of unique authors per year. **C**. Phage research experienced a steep increase in recent popularity. **D**. TAELNs, ZFNs, and CRISPR are three competing gene editing technologies, with clear field preferences. The popularity of each field is clearly visualized by author count with peaks and valleys expectedly following the development of each new technique.

## Discussion

Understanding field size and organization is essential to effectively organizing our scientific workforce. There is some evidence that scientists favor working on specific topics, which raises the concern chasing “hot-topics” could neglect essential basic science research^10^. We hoped to develop a measurement of field size which could help quantify these differences and determine how and where our scientific work-effort is distributed.

We found that counting unique authors provided an estimate of the size and activity of a given field. The overall number of papers published in a field each year trended strongly with the total number of unique authors across many, but not all, of the analyzed fields. Importantly, the use of unique authors in a field differs from merely counting total publications in that it offers additional dimensions of analysis. Publication counts reflect research output of a field as a whole but understanding how that output is distributed across different laboratories and working environments gives a more granular image of the research landscape. Diversity of ideas and research groups is essential to creative science and fostering new ideas whereas fields dominated by fewer smaller lab groups may find themselves stymied from dogmatic hierarchies.

Measuring unique authors also avoids certain pitfalls inherent to a paper number driven analyses. First, it allows us to control for differences in output between individual authors and fields. Since we are looking to measure effort and workforce rather than productivity, a single author who outputs 10 articles per year should be weighed the same as an author who outputs 15. Conversely, a paper with 10 authors implies a larger workforce behind it than a paper with 2. Thus, quantifying effort and personnel according to papers does not allow for granularity in terms of personnel and may explain why we sometimes observed differences in field size when comparing total paper counts to unique author counts.

NIH RePORTER and CDC Required Reporting data is limited in the context of these diseases but we were able to identify a slight trend of increasing funding per author. When comparing the size of a given field to the public health burden of its respective disease, we noticed a positive correlation across time and irrespective of whether the focus was on global cases or limited to the United States. We observed a similar trend when comparing the WHO priority list of fungal pathogens, noting that critical fungal pathogens on average had a larger proportion of the workforce, followed by high priority, followed by medium priority. This analysis is reassuring that the allocated labor force is efficiently focusing on microbes with high disease burden while not neglecting microbes with lower burden diseases. This trend persisted even when comparing fungal diseases to more globally significant human diseases such as HIV, malaria, and tuberculosis. These larger fields experienced similar growth over time, but the distribution of funding remained similar to the fungal fields, further reassuring us that resources and workforce distributes efficiently between fields of research rather than disproportionately flocking to perceived ‘hot-topic’ research.

Our analysis also provides insight into the growth and decline of fields over time. For example, the use and popularity of the *S. cerevisiae* system as a model for eukaryotic cell biology rose rapidly in the 1970s-80s and then stabilized, presumably as mammalian cell systems matured to allow comparable types of experimentation. In 1990 the estimated size of the four model organisms was *S. cerevisiae* > *D. melanogaste*r > *C. elegans* while zebra fish had not yet emerged as a major model organism. Three decades later, there are more authors associated with zebra fish than the other model organisms and *S. cerevisiae* has moved to second place. We interpret this sequence as reflecting the fact that for eukaryotic cell biology, molecular techniques for studying deep questions were available first in yeast, then invertebrate flies, and most recently in the vertebrate zebra fish. In virology, simian vacuolating virus 40 (SV40), a polyoma virus that can cause tumors was a major early experimental system used to understand how viruses caused cancer, but its popularity has declined as investigators have moved to other systems. Poliovirus was a major medical problem before the introduction of effective vaccines in the 1950s and the number of authors associated with papers on this virus remained relatively stable until recent years when the virus resurfaced with new strains and a decline in vaccination in some regions. Similarly, smallpox was a scourge in the past but was eradicated in 1977 and followed by steady decline around 1990, then a resurgence of interest with concerns about its potential as a biological weapon and increased interest given human pathogenic potential of other poxviruses. Hence, for both model systems and three major viruses the trends in author numbers can be associated with historical developments in their fields.

Our analysis has several caveats and limitations. First, there is not complete standardization of author name format in PubMed, meaning that the same author may appear and be counted twice in our analysis if, for example, their name appears as LAST FIRST and LAST FIRST-INITIAL on separate publications. We were able to identify and correct this issue for authors we personally knew, but it would be impossible to parse out initials from more common names (ex. Smith). With the increasing prevalence of ORCIDs this problem may be fixed in the future. We also caution that field size estimated from author numbers is likely to be an upper ceiling estimate since not everyone who authors a paper in an area is necessarily doing research in the area in question. For example, the large increase in investigators authoring phage and CRISPR papers likely reflects the usefulness of those systems for a variety of high throughput experiments, rather than an increase in the numbers of individuals working on phage biology or CRISPR machinery.

In summary, we propose a bibliometric method for estimating the size of fields and apply it to microbiology and several subfields. When applied to the subfield of medical mycology the results show differences in the size of fields that correlate with the medical importance of a particular fungus. We are hopeful that this approach provides a useful tool for sociologists of science and policy makers for studying the structure of scientific fields.

## Acknowledgments

This work is supported by NIH grants 5R01HL059842-23 and 1R01AI171093-01A1.

---

**Supplemental Figure 1.**
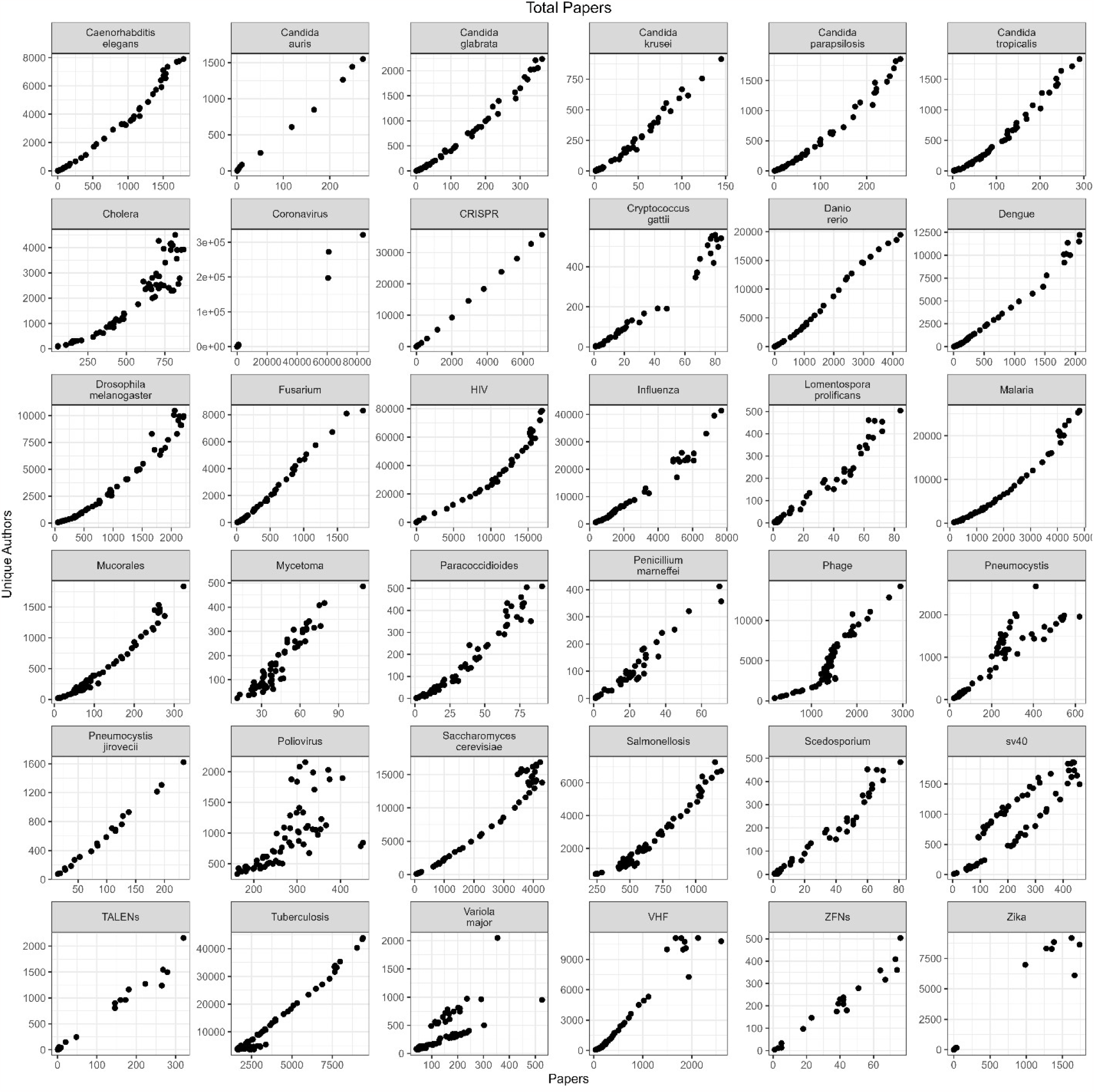
Size of field comparison between author and paper count. We compared paper count to unique author count as estimates for size of fields. Overall, the two correlate well, suggesting that paper could be used as a broad surrogate measure of field size.

